# Crumble: reference free lossy compression of sequence quality values

**DOI:** 10.1101/243030

**Authors:** James K Bonfield, Shane A McCarthy, Richard Durbin

## Abstract

**Motivation:** The bulk of space taken up by NGS sequencing CRAM files consists of per-base quality values. Most of these are unnecessary for variant calling, offering an opportunity for space saving.

**Results:** On the CHM1+CHM13 test set, a 17 fold reduction in quality storage can be achieved while maintaining variant calling accuracy.

**Availability:** Crumble is OpenSource and can be obtained from https://github.com/jkbonfield/crumble.

**Supplementary information:** Supplementary data are available.

## 1 Introduction

The rapid reduction of costs for genome sequencing (Wetterstrand, 2016) has led to a corresponding growth in storage costs, far outstripping Moore’s Law for CPU and Kryder’s Law for storage. This has led to considerable research into DNA sequence data compression (Numanagić *et al*., 2016).

The most significant component in data storage cost is the per-nucleotide confidence values, which carry information about the likelihood of each base call being in error. The original CRAM proposal (Fritz *et al*., 2011) introduced the term ‘quality budget’ for lossy compression. Given a fixed amount of storage we can decide how to spend this budget, either by uniform degradation of all qualities or more targeted fidelity in important regions only. How to target this has been the focus of lossy compression research, with two main strategies: “horizontal” and “vertical”.

“Horizontal” compression smooths qualities along each sequence in turn, as implemented in libCSAM (Cánovas *et al*., 2014), QVZ (Malysa *et al*., 2015) and FaStore (Roguski *et al*., 2017) or via quantisation (Illumina, 2014). This type of compression can be applied before alignment and is entirely reference free.

“Vertical” compression takes a slice through an aligned dataset in the SAM format (Li *et al*., 2009) to determine which qualities to keep and which to discard, as used in CALQ (Voges *et al*., 2017), or via hashing techniques on unaligned data in Leon (Benoit *et al*., 2015) and GeneCodeq (Greenfield *et al*., 2016). Traditional loss measures, such as mean squared error, will appear very high, but these tools focus on minimising the changes in post-processed data (variant calling).

We present Crumble as a mixture of both horizontal and vertical compression. It operates on coordinate sorted aligned SAM, BAM or CRAM files. While this approach does not explicitly use a reference, the sequence aligner does, which may result in some reference bias.

## 2 Methods

A variant caller evaluates the sequence base calls overlapping each genome locus along with their associated qualities to determine whether that site represents a variant. Irrespective of whether the call is a variant, if the same call is made with comparable confidence both with and without sequence quality values present then it can be concluded that the qualities are not necessary in that column.

This requires running the variant caller twice to assess the change, but if limited to sites with high confidence calls the need for a second test can be avoided. We implemented a fast, but naïve, caller derived from Gap5’s consensus algorithm (Bonfield and Whitwham, 2010). This is a pure pileup-column oriented approach that treats the lack of a base (deletion) as a 5th base type (“*”) and then identifies the most likely homozygous or heterozygous combination of bases that match the observed base calls, confidence values and mapping qualities. Thus it assumes a single individual diploid sample and is not tuned to work with somatic variants. This is further modified by reducing the confidence for the consensus by the bases which do not match the hypothesis, thus producing a deliberately pessimistic caller. The aim is not to have a built-in high-quality caller, but to preserve quality values if any downstream variant caller may be uncertain while retaining independence from any standard tool.

Even when deemed unnecessary, qualities cannot be entirely discarded as tools expect them to exist. By replacing the qualities for bases that agree with a confident consensus call with constant high values, the entropy of the quality signal is reduced. Quality values for bases that disagree with a confident consensus call may optionally be set to a constant low value, heavily quantised, or left intact.

There are sites where any variant caller may incorrectly give the wrong call with high confidence. Furthermore the reference itself may be incorrect and a subsequent realignment to an updated reference may change read locations and alignment strings. We do not wish to replace qualities in such regions. We therefore have a set of heuristics to try to find potentially unreliable calls and retain verbatim the confidence values for these locations and surrounding bases depending on sequence context. Similarly there may be places where an entire read needs to have qualities retained as there is evidence for it being misplaced or being part of a large structural rearrangement.

The heuristics used in Crumble to identify where confidence values should be retained vary by compression level requested, but include:

- **Concordant soft clipping**: many reads having soft clipped bases at the same site often indicates a large insertion (absent in the reference) or contamination.
- **Excessive depth**: possible contamination or collapsed repeat. Variant calls often appear unusually good in such data, even when wrong.
- **Low mapping quality**: possibly caused through poor reference. We optionally can also store quality values for the reads with high mapping quality that colocate with many low mapping quality reads.
- **Unexpected number of variants**: we assume data from a single diploid sample with at most two alleles at each locus. More than two alleles implies misaligned data, duplication or contamination.
- **Low quality variant calls**: typically a single base locus where the consensus is unclear.
- **Proximity to short tandem repeats**: alignments are often poor in such regions, especially if indels are present, leading to bases occurring in the wrong pileup column.

Finally for the quality values that we deem necessary to keep, we optionally provide horizontal compression via the P-block algorithm from CSAM. This is most useful on older Illumina data sets with over 40 distinct levels of quality values.

The nature of the Crumble algorithm makes it amenable to streaming and it does not require large amounts of memory to operate.

## 3 Results

Analysis of how quality compression affects variant calling was performed on Syndip (Li *et al*., 2017), an Illumina sequenced library artificially constructed from the haploid cell lines CHM1 and CHM13, with an associated high quality truth set based on two PacBio assemblies (Schneider *et al*., 2017). Compared to the Genome in a Bottle (GIAB) or Platinum Genomes (PlatGen) data sets this has a considerably larger set of tricky indels in the truth set, giving SNP false positive rates 5-10 times higher (Li *et al*., 2017) than on GIAB or PlatGen truth sets. While Syndip still requires a list of regions to exclude, the total number of excluded non-N reference bases is 40% fewer than GIAB 3.3.2. By restricting analysis to solely the regions within Syndip and not within GIAB we observe 65% of chromosome 1 false positives occur within this region, but crumble still shows good performance (see Supplementary Data).

The input BAM file (ERR1341796) had previously been created with GATK best practices including IndelRealigner and Base Quality Score Recalibration steps. To test the impact on raw variant calling, we ran GATK HaplotypeCaller (Poplin *et al*., 2017), Bcftools (Li, 2011) and Freebayes (Garrison and Marth, 2012), filtering to calls of quality 30 or above, without use of GATK Variant Quality Score Recalibration. As a baseline we compare Crumble to the original lossless results and against a single fixed quality value. This latter test demonstrates that quality values are important, but we only need a small quality budget to achieve comparable results to lossless compression. Indeed, we observe that vertical quality score compression can marginally improve variant calling by standard callers, as has been noted previously in the QVZ (Malysa *et al*., 2015) and Leon (Benoit *et al*., 2015) papers.

Table 1 shows the GATK lossless results on the CHM pair along with the changes caused by lossy compression using a variety of Crumble options on both the full Syndip data and a low coverage subset. We chose the minimal compression level, an expected maximum compression level and a set of manually tuned parameters optimised for this data set. The manual tuning traded false positives and false negatives in an attempt to get a call set comparable or better than the original in all regards. It is unknown if the tuned parameters are appropriate for all data sets. More complete comparisons including against other tools are available in the online Supplementary Data.

**Table 1.** Effect of lossy quality compression on 50x and 15x Syndip data using GATK HaplotypeCaller

| Category | Original | Original F | Crumble-1 | Crumble-1 F | Crumble-9p8 | Crumble-9p8 F | Crumble* | Crumble* F |
| --- | --- | --- | --- | --- | --- | --- | --- | --- |
| 50x Qual size (MB) | 4107 | - | 614 | - | 235 | - | 229 | - |
| 50x SNP False Positive | 6226 | 2968 | -359 | -79 | -251 | -67 | -526 | -181 |
| 50x SNP False Negative | 4648 | 7625 | 0 | -53 | -25 | -184 | +41 | -123 |
| 50x Indel False Positive | 3965 | 3649 | -7 | -41 | +19 | +9 | -35 | -32 |
| 50x Indel False Negative | 7881 | 7972 | +7 | +11 | -103 | -82 | -93 | -72 |
| 15x Qual size (MB) | 1211 | - | 260 | - | 77 | - | 72 | - |
| 15x SNP False Positive | 4798 | 2517 | -10 | +63 | +347 | +225 | -359 | -29 |
| 15x SNP False Negative | 14985 | 27761 | -205 | -297 | -3027 | -4608 | -1866 | -2865 |
| 15x Indel False Positive | 2781 | 2521 | +2 | -14 | +109 | +60 | +53 | +26 |
| 15x Indel False Negative | 13136 | 13925 | -8 | +5 | -484 | -427 | -444 | -410 |

Comparison of unfiltered and filtered (marked with “F”) calls on the Syndip truth set. GATK filtering rules are listed in the Supplementary Data. Crumble\* refers to parameters optimised for this data set: “crumble -9p8 -u30 -Q60 -D100”. The false positive/negative values of the GATK calls on the crumbled data set are shown relative to their respective GATK called lossless dataset. The truth set for chromosome 1 has 269,655 SNPs and 46,036 indels, counting multi-allelic sites once per allele. The quality sizes are absolute for all files.

On the original BAM file with ∼50x coverage we observed a 17 fold reduction in the size of CRAM compressed quality values, while achieving a 6% drop in filtered SNP false positive rate (higher precision) and 2% drop in false negative rates (higher recall). Indels also see a 1% improvement in both measures. At a sub-sampled 15x coverage we see a 1% drop in filtered SNP false positive rates and a 10% reduction in SNP false negatives. Indel calls were more comparable, with 1% higher false positives and 3% lower false negatives.

It is likely these gains to both SNP precision and recall only apply to data coming from a single individual, but they demonstrate the efficacy of lossy quality compression.

## 4 Conclusion

We have demonstrated that Crumble, when combined with CRAM, can greatly reduce file storage costs while having a minimal, if not beneficial, impact on variant calling accuracy of individual samples. For maximum compression Crumble also permits discarding read identifiers and some auxiliary tags, typically yielding files in the size of 5-10Gb for a 30x whole genome processed with Crumble −9p8. Using this across a variety of BAM and CRAM files Crumble gave an overall file size reduction from 3 to 7.8 fold (details in Supplementary Data).

Crumble is designed to operate on a single sample file. For multiple samples, it is best to apply Crumble to each sample independently, produce gVCF, and then jointly call from those. Note Crumble is explicitly designed to operate on diploid data, so it is not appropriate for use on sequence from cancer or other samples with subclonal genetic structure.

## Acknowledgements

We would like to thank Yasin Memari for help testing and evaluating an earlier version of the program.

