## Supplementary Materials for "Crumble: reference free lossy compression of sequence quality values"

### Crumble: supplementary material

March 22, 2018

#### 1 Introduction

Crumble does not compress quality values itself, rather it replaces quality values in a SAM/BAM/CRAM file with different qualities which compress better in standard tools. If the distribution of quality value usage becomes more extreme, the entropy decreases and compression ratios increase.

This means that existing software pipelines continue to work on crumbled data. However it also means some file formats gain more from Crumble than others.

#### 2 Software versions and git commit hashes

| | | | |
|---|---|---|---|
| Crumble | 0.8 | 996341e | `https://github.com/jkbonfield/crumble` |
| GATK | 3.7 | | `https://software.broadinstitute.org/gatk` |
| CALQ | 1.0.0 | 5b2ba4c | `https://github.com/voges/calq` |
| Bcftools | 1.6-7 | b7b502e | `https://github.com/samtools/bcftools` |
| Freebayes | 1.1.0-46 | 8d2b3a0 | `https://github.com/ekg/freebayes` |
| QVZ2 | 0.1-24 | 70e5926 | `https://github.com/mikelhernaez/qvz2` |
| VT | 0.5772 | 6686b5c | `https://github.com/atks/vt` |

#### 3 Evaluation pipeline

GATK HaplotypeCaller, Bcftools and Freebayes are used without a set of known variants and without application of GATK Variant Quality Score Recalibration (VQSR). This is to demonstrate the raw calling power without attempts to rescue mistakes via known variants and to judge likely performance on new organisms. Command line arguments used were:

```
java -Xmx4g -jar GenomeAnalysisTK.jar -T HaplotypeCaller -R $human_ref \
  -L 1 --genotyping_mode DISCOVERY -stand_call_conf 10 \
  -I $prefix.bam -o $prefix.gatk.vcf

freebayes -f $human_ref $prefix.bam > $prefix.freebayes.vcf

bcftools mpileup -f $human_ref $prefix.bam  | \
  bcftools call -vm - > $prefix.bcftools.vcf
```

Truth sets are downloaded from Heng Li's CHM-eval release:
`https://github.com/lh3/CHM-eval/releases/download/v0.2/CHM-evalkit-20161018.tar`

Comparison of VCF call and truth sets is made after normalising variant coordinates and splitting multi-allelic sites and MNPs into separate vcf records, followed by region filtering using the inclusion / exclusion bed files in the CHM-eval release kit. The effect of these may mean that some compound variants can yield both a match and a mismatch, for example calling a homozygous mutation as heterozygous, but it makes comparisons between tools easier. These operations are performed with bcftools and vt:

```
bcftools norm -m -both -t $region -f $href $v 2>/dev/null | \
  vt decompose_blocksub - | \
  bcftools view -T ^$exclude.bed | bcftools view -T $include.bed > $v.norm.vcf
```

The normalised / filtered files are then compared with "bcftools isec" to count the shared variants between truth and call sets and those occurring only in one file:

```
bcftools isec -c both -p $call.isec $truth.norm.vcf.gz $call.norm.vcf.gz
```

This is a relatively strict definition of identity, meaning that the variant must occur at both the same site and be the same call. The "isec" command produces 4 VCF files in the `$call.isec` directory:

`0000.vcf`: private to truth.norm.vcf (false negatives)
`0001.vcf`: private to call.norm.vcf (false positives)
`0002.vcf`: records from truth.norm.vcf, shared by both files (correct calls)
`0003.vcf`: records from call.norm.vcf, shared by both files (correct calls)

By counting the VCF records in each file we observe the recall and precision. The files can be filtered by quality and type using "bcftools view", for example:

```
FN_SNP=‘bcftools view -H -i "TYPE=’snp’ && QUAL >= 30" $call.isec/0001.vcf | wc -l‘
```

More aggressive filtering was also applied based on the recommended practices from each tool, where available. The following are exclusion filter rules, applied using 'bcftools view -e $filter'. We also applied a simple over-depth filter too, of DP>90 for the full 50x sample and DP>30 for the 15x sample.

- **GATK HaplotypeCaller**
  https://software.broadinstitute.org/gatk/documentation/article.php?id=3225

  **SNP**: `QUAL < $qual || QD < 2 || FS > 60 || MQ < 40 || SOR > 3 || MQRankSum < -12.5 || ReadPosRankSum < -8 || DP > $DP`

  **Indel**: `QUAL < $qual || QD < 2 || FS > 200 || ReadPosRankSum < -20 || DP > $DP`

- **Bcftools** (No quality filtering for indels)

  **SNP**: `QUAL < $qual || DP > $DP`

  **Indel**: `IDV < 3 || IMF < 0.03 || DP > $DP`

- **Freebayes**
  https://wiki.uiowa.edu/download/attachments/145192256/erik%20garrison%20-%20iowa%20talk%202.pdf?api=v2

  **SNP / Indel**: `QUAL < $qual || SAF <= 0 || SAR <= 0 || RPR <= 1 || RPL <= 1 || DP > $DP`

Note that due to some variants being compound, it is possible for a single VCF record to contain the correct variant while also containing either a false positive or false negative.

It is also noted that the normalisation step is not always perfect and we cannot compute whether a compound insertion and deletion is identical to a series of SNPs. Hence some of the reported numbers of false positives / negatives may be pessimistic. However we do not believe the results are biased in favour of any specific method of quality reduction.

### 4 Results

The original BAM input file was chromosome 1 of CHM1_CHM13_2.bam, from ERR1341796 with depth ~50x. We also subsampled this to evaluate performance on a ~15x data set, where quality values become much more important.

The first assessment we do is to evaluate the baseline of lossless quality values, followed by no quality values (using a fixed score) to demonstrate the impact that having any quality has. Subsequent tests evaluate quality quantisation, Crumble, Calq and QVZ2. We test variant calling precision and recall using GATK HaplotypeCaller, Bcftools and Freebayes.

Tables below show the number of true positives (TP), false positives (FP) and false negatives (FN) for all variants, after filtering by quality, and with a more complete filtering by quality, depth and per-tool recommended rules.

For our tables we use variant quality 30 in our filters, but variant callers calibrate quality values differently and the trade off between precision and recall may alter at a different quality threshold. To get a better comparison between tools and the effect that variant quality filtering has on each tool we plot the true positive vs false positive rates as a line, with points produced by varying the quality filter to values 10, 15, 20, 25, 30, 40, 50, 75 and 100. Points closer to the top-left of the graph represent a better result with fewer false positive and/or false negative calls. Each tool is graphed with and without the additional filtering steps listed in the introduction.

#### 4.1 Original / Quantised, Chromosome 1

We first present the baseline original quality values for Chromosome 1 of the download BAM file along with no quality values using a fixed quality of 25, and simple binary quantisation with qualities 4 and 28. The reason to consider these one and two value quantisations is to provide a baseline for more targeted approaches.

We count the total number of bases in chromosome 1 alignments along with the expected number of base call errors according to their quality values. For example, if we observe 1,000 bases with phred quality 20 then we expect approximately 10 will be erroneous as quality 20 (assuming a correct BQSR recalibration) indicates a 1 in 100 error rate. For the full 50x data on chromosome 1 this gives 12,239,915,644 bases with an estimated 599,904,677 errors, yielding an amortised average quality score of 13.1. Unfortunately using this gives no calls with GATK and a large number of false negatives using bcftools and freebayes. So instead we chose an arbitrary quality value of 25 as a means to evaluate quality-less performance.

For the binary quantisation, we observe a dip in the quality frequency distribution between 16 and 20, so we split the distribution into bases with quality >= 20 and those below. By similar counting these lead to amortised base quality scores of 4 and 28 for the two bins, which unlike single quality 13 does work well for all three tools.

The binary quantisation using values 4 and 28 has minimal impact on bcftools and freebayes recall and accuracy. With GATK it also has minimal impact on the 15x data, but with the 50x it has a small negative impact.

All three callers perform poorly with the unary quality 25, with significant increases in either false positives (Bcftools, Freebayes) or false negatives (GATK). Thus we establish that some degree of quality value separation is important for calling accuracy, even at 50 fold coverage. While using a unary quality would effectively remove all storage requirements for quality values, the binary quantisation compresses quality value storage by a factor of 7.6.

#### GATK HaplotypeCaller

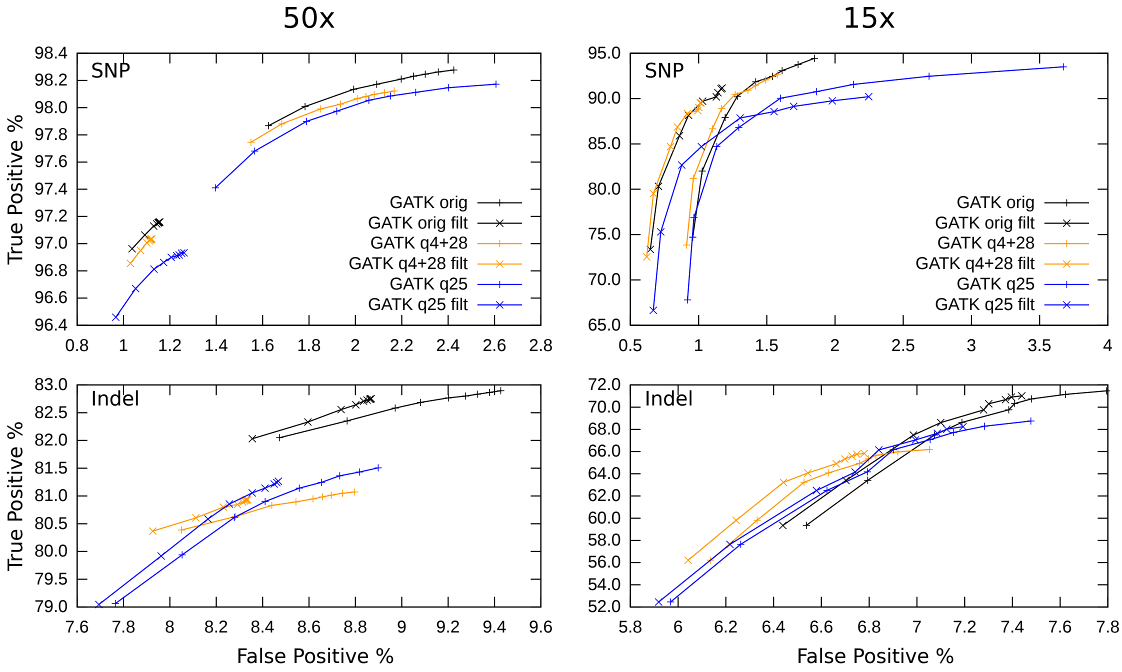


Figure 1: *True Positive vs False Negative rates of GATK HaplotypeCaller on the original qualities vs binary and unary quantisation.*

These show that having some fidelity of quality values is beneficial, as the fixed value of 25 does not compare well to the original. Binary quantisation to 4 and 28 has a negative impact on GATK at high depth, but minimal change on the shallow data set.

Tables with the actual counts of true positives, false positives and false negatives are shown below.

Table 1: GATK HC: 50x Original

| Type | | Q>0 | Q>=30 | Filtered |
|---|---|---|---|---|
| SNP | TP | 265007 | 264828 | 261977 |
| SNP | FP | 6585 | 5950 | 3047 |
| SNP | FN | 4648 | 4827 | 7678 |
| InDel | TP | 38162 | 38103 | 38075 |
| InDel | FP | 3972 | 3861 | 3690 |
| InDel | FN | 7874 | 7933 | 7961 |

**CRAM qual size 4,106,563,351**

Table 2: GATK HC: 15x Original

| Type | | Q>0 | Q>=30 | Filtered |
|---|---|---|---|---|
| SNP | TP | 254670 | 247683 | 241894 |
| SNP | FP | 4798 | 3564 | 2517 |
| SNP | FN | 14985 | 21972 | 27761 |
| InDel | TP | 32900 | 32117 | 32111 |
| InDel | FP | 2781 | 2561 | 2521 |
| InDel | FN | 13136 | 13919 | 13925 |

**CRAM qual size 1,211,486,517**

Table 3: GATK HC: 50x Qual 4 + 28

| Type | | Q>0 | Q>=30 | Filtered |
|---|---|---|---|---|
| SNP | TP | 264592 | 264442 | 261645 |
| SNP | FP | 5861 | 5418 | 2950 |
| SNP | FN | 5063 | 5213 | 8010 |
| InDel | TP | 37322 | 37265 | 37238 |
| InDel | FP | 3600 | 3514 | 3377 |
| InDel | FN | 8714 | 8771 | 8798 |

**CRAM qual size 539,249,433**

Table 4: GATK HC: 15x Qual 4 + 28

| Type | | Q>0 | Q>=30 | Filtered |
|---|---|---|---|---|
| SNP | TP | 249779 | 243924 | 238273 |
| SNP | FP | 4000 | 3132 | 2206 |
| SNP | FN | 19876 | 25731 | 31382 |
| InDel | TP | 30470 | 29891 | 29884 |
| InDel | FP | 2312 | 2167 | 2133 |
| InDel | FN | 15566 | 16145 | 16152 |

**CRAM qual size 159,104,061**

Table 5: GATK HC: 50x Qual 25

| Type | | Q>0 | Q>=30 | Filtered |
|---|---|---|---|---|
| SNP | TP | 264727 | 264408 | 261295 |
| SNP | FP | 7085 | 5556 | 3189 |
| SNP | FN | 4928 | 5247 | 8360 |
| InDel | TP | 37522 | 37354 | 37315 |
| InDel | FP | 3665 | 3496 | 3402 |
| InDel | FN | 8514 | 8682 | 8721 |

**CRAM qual size 756,507**

Table 6: GATK HC: 15x Qual 25

| Type | | Q>0 | Q>=30 | Filtered |
|---|---|---|---|---|
| SNP | TP | 252113 | 242781 | 236923 |
| SNP | FP | 9614 | 3946 | 3132 |
| SNP | FN | 17542 | 26874 | 32732 |
| InDel | TP | 31651 | 30461 | 30451 |
| InDel | FP | 2558 | 2258 | 2236 |
| InDel | FN | 14385 | 15575 | 15585 |

**CRAM qual size 223,176**

#### Bcftools

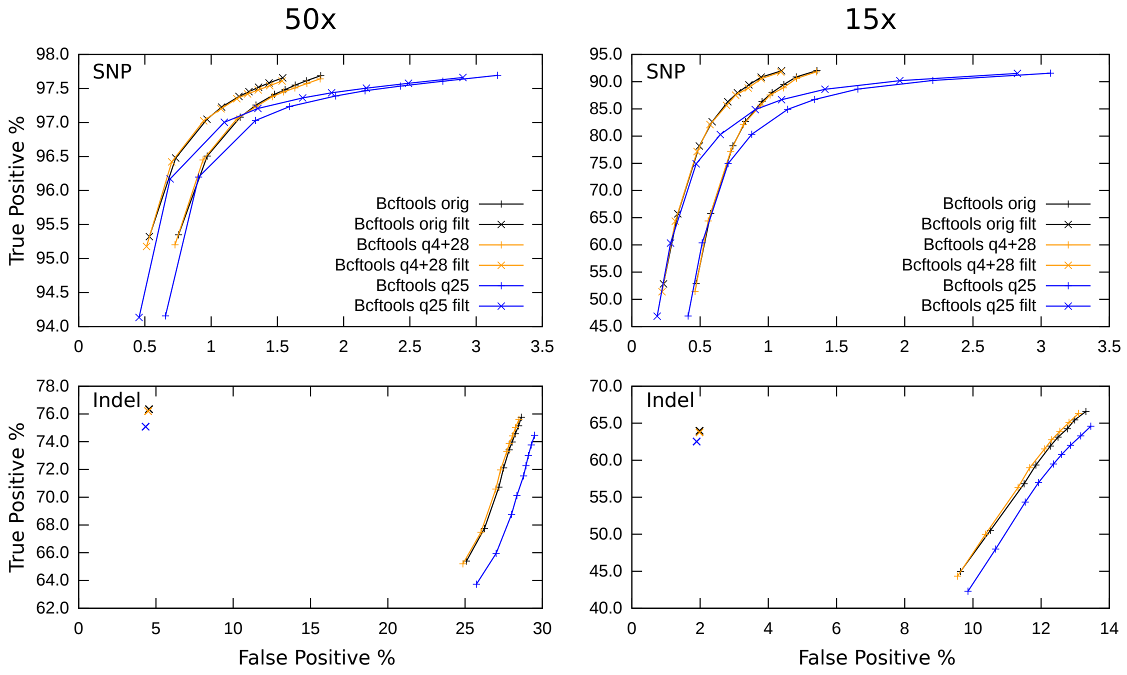


Figure 2: *True Positive vs False Negative rates of Bcftools on the original qualities vs binary and unary quantisation.*

As with GATK HaplotypeCaller, Bcftools is harmed by having no quality values. However the lines showing binary binned (4 and 28) qualities are nearly superimposed on top of the lossless quality calls, at some points being marginally improved by the binning process.

Note the bcftools indel filtering doesn't use quality values, hence these come out as a single point.

Table 7: Bcftools: 50x Original

| Type | | Q>0 | Q>=30 | Filtered |
|---|---|---|---|---|
| SNP | TP | 263750 | 262682 | 262599 |
| SNP | FP | 5493 | 3942 | 3216 |
| SNP | FN | 5905 | 6973 | 7056 |
| InDel | TP | 35434 | 33799 | 35143 |
| InDel | FP | 14490 | 13048 | 1678 |
| InDel | FN | 10602 | 12237 | 10893 |

**CRAM qual size 4,106,563,351**

Table 8: Bcftools: 15x Original

| Type | | Q>0 | Q>=30 | Filtered |
|---|---|---|---|---|
| SNP | TP | 253194 | 232858 | 232734 |
| SNP | FP | 4763 | 2243 | 1648 |
| SNP | FN | 16461 | 36797 | 36921 |
| InDel | TP | 31820 | 28502 | 29450 |
| InDel | FP | 5198 | 3985 | 596 |
| InDel | FN | 14216 | 17534 | 16586 |

**CRAM qual size 1,211,486,517**

Table 9: Bcftools: 50x Qual 4 + 28

| **Type** | | **Q>0** | **Q>=30** | **Filtered** |
|---|---|---|---|---|
| SNP | TP | 263644 | 262590 | 262507 |
| SNP | FP | 5521 | 3895 | 3171 |
| SNP | FN | 6011 | 7065 | 7148 |
| InDel | TP | 35364 | 33737 | 35080 |
| InDel | FP | 14360 | 12923 | 1652 |
| InDel | FN | 10672 | 12299 | 10956 |

**CRAM qual size 539,249,433**

Table 10: Bcftools: 15x Qual 4 + 28

| **Type** | | **Q>0** | **Q>=30** | **Filtered** |
|---|---|---|---|---|
| SNP | TP | 252813 | 231041 | 230917 |
| SNP | FP | 4945 | 2203 | 1613 |
| SNP | FN | 16842 | 38614 | 38738 |
| InDel | TP | 31672 | 28325 | 29354 |
| InDel | FP | 5102 | 3900 | 594 |
| InDel | FN | 14364 | 17711 | 16682 |

**CRAM qual size 159,104,061**

Table 11: Bcftools: 50x Qual 25

| **Type** | | **Q>0** | **Q>=30** | **Filtered** |
|---|---|---|---|---|
| SNP | TP | 263813 | 262617 | 262539 |
| SNP | FP | 11444 | 5196 | 4515 |
| SNP | FN | 5842 | 7038 | 7116 |
| InDel | TP | 34831 | 32932 | 34564 |
| InDel | FP | 14830 | 13308 | 1567 |
| InDel | FN | 11205 | 13104 | 11472 |

**CRAM qual size 756,507**

Table 12: Bcftools: 15x Qual 25

| **Type** | | **Q>0** | **Q>=30** | **Filtered** |
|---|---|---|---|---|
| SNP | TP | 252531 | 228972 | 228851 |
| SNP | FP | 17447 | 2646 | 2088 |
| SNP | FN | 17124 | 40683 | 40804 |
| InDel | TP | 30978 | 27389 | 28782 |
| InDel | FP | 5128 | 3863 | 557 |
| InDel | FN | 15058 | 18647 | 17254 |

**CRAM qual size 223,176**

##### Freebayes

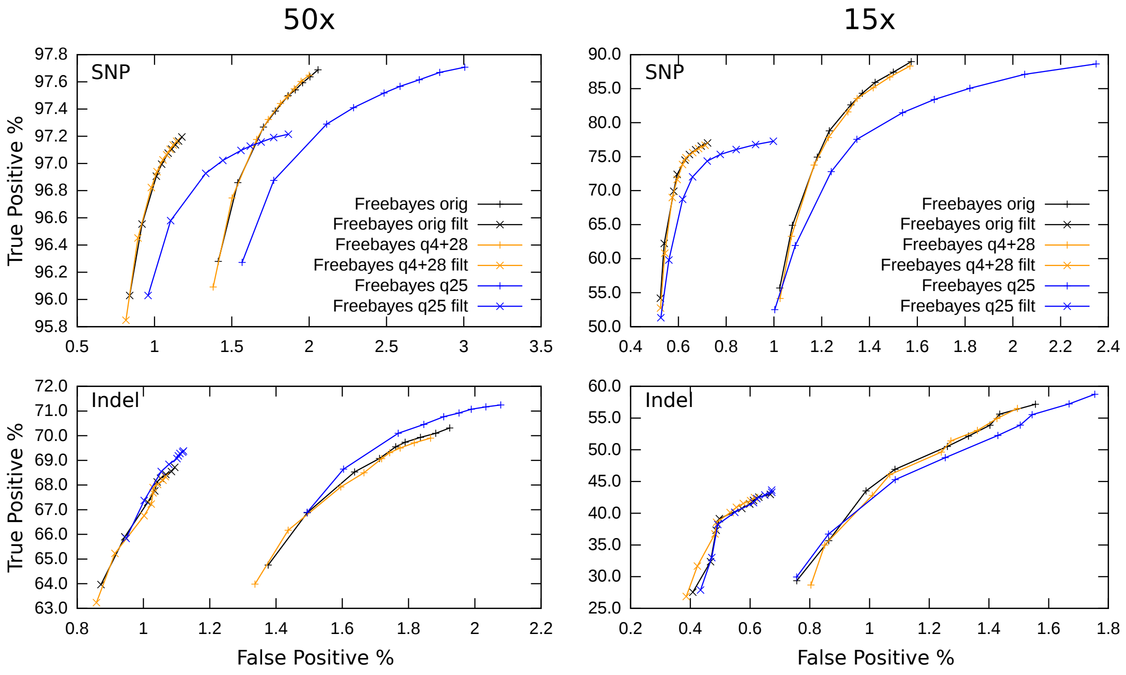


Figure 3: *True Positive vs False Negative rates of Freebayes on the original qualities vs binary and unary quantisation.*

As with Bcftools, fixed quality is harmful, but again we see binary quantisation having either no effect or a small benefit.

Table 13: Freebayes: 50x Original

| Type | | Q>0 | Q>=30 | Filtered |
|---|---|---|---|---|
| SNP | TP | 264313 | 262909 | 261769 |
| SNP | FP | 6018 | 4994 | 2880 |
| SNP | FN | 5342 | 6746 | 7886 |
| InDel | TP | 32756 | 32018 | 31362 |
| InDel | FP | 675 | 574 | 330 |
| InDel | FN | 13280 | 14018 | 14674 |

**CRAM qual size 4,106,563,351**

Table 14: Freebayes: 15x Original

| Type | | Q>0 | Q>=30 | Filtered |
|---|---|---|---|---|
| SNP | TP | 258868 | 222751 | 200892 |
| SNP | FP | 4994 | 2984 | 1269 |
| SNP | FN | 10787 | 46904 | 68763 |
| InDel | TP | 30122 | 23257 | 18760 |
| InDel | FP | 535 | 297 | 108 |
| InDel | FN | 15914 | 22779 | 27276 |

**CRAM qual size 1,211,486,517**

Table 15: Freebayes: 50x Qual 4 + 28

| Type | | Q>0 | Q>=30 | Filtered |
|---|---|---|---|---|
| SNP | TP | 264310 | 262753 | 261637 |
| SNP | FP | 5863 | 4856 | 2789 |
| SNP | FN | 5345 | 6902 | 8018 |
| InDel | TP | 32687 | 31795 | 31159 |
| InDel | FP | 654 | 556 | 324 |
| InDel | FN | 13349 | 14241 | 14877 |

**CRAM qual size 539,249,433**

Table 16: Freebayes: 15x Qual 4 + 28

| Type | | Q>0 | Q>=30 | Filtered |
|---|---|---|---|---|
| SNP | TP | 258878 | 219981 | 199141 |
| SNP | FP | 5018 | 2919 | 1236 |
| SNP | FN | 10777 | 49674 | 70514 |
| InDel | TP | 30098 | 22849 | 18462 |
| InDel | FP | 532 | 287 | 99 |
| InDel | FN | 15938 | 23187 | 27574 |

**CRAM qual size 159,104,061**

Table 17: Freebayes: 50x Qual 25

| Type | | Q>0 | Q>=30 | Filtered |
|---|---|---|---|---|
| SNP | TP | 264306 | 262960 | 261822 |
| SNP | FP | 9670 | 6698 | 4147 |
| SNP | FN | 5349 | 6695 | 7833 |
| InDel | TP | 32964 | 32578 | 31797 |
| InDel | FP | 739 | 633 | 354 |
| InDel | FN | 13072 | 13458 | 14239 |

**CRAM qual size 756,507**

Table 18: Freebayes: 15x Qual 25

| Type | | Q>0 | Q>=30 | Filtered |
|---|---|---|---|---|
| SNP | TP | 258860 | 219683 | 200481 |
| SNP | FP | 11610 | 3433 | 1455 |
| SNP | FN | 10795 | 49972 | 69174 |
| InDel | TP | 30255 | 24061 | 19185 |
| InDel | FP | 631 | 349 | 118 |
| InDel | FN | 15781 | 21975 | 26851 |

**CRAM qual size 223,176**

##### Tool Comparisons

Given the above analysis, we are also able to do a side by side comparison between GATK HaplotypeCaller, Bcftools and Freebayes results on both 50x and 15x data sets. Such an analysis is not the primary focus of this paper, but given we have the data available it is an interesting diversion.

Missing from these figures is the usefulness of output. In order to compare between tools and get a constant total number of variants we have split all multi-allelic sites and MNPs into individual records, as this permits Freebayes haplotype calls to be compared against bcftools and GATK HaplotypeCaller, however in doing so it removes one of the strengths of Freebayes in that neighbouring mutations are phased. It should be noted this is purely a snapshot of one single individual with two alleles in even proportion, so we do not encourage any broader conclusions to be made. Also note that regardless of the tool used for calling, the data has previously been passed through GATK BQSR (base quality score recalibration).

On this data set we observe that each tool occupies its own distinct space in the accuracy (true positives) vs recall (false negatives) graph for SNP calling, meaning that each tool has its own strengths.

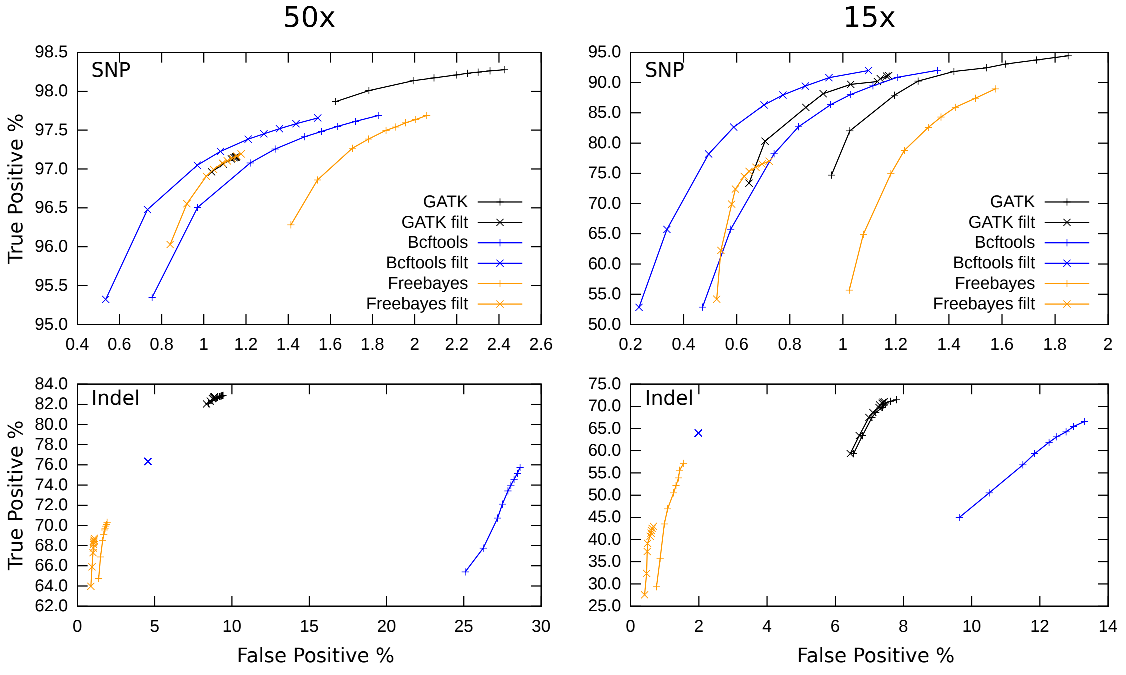


Figure 4: *A summary of True Positive vs False Negative rates of GATK HaplotypeCaller, Bcftools and Freebayes at multiple quality thresholds, with and without filtering.*

#### 4.2 Crumble

Crumble was tested with minimum (-1), maximum (-9p8) and custom optimised (-9p8 -u30 -Q60 -D100) parameters. The compression level (1 to 9) controls a larger set of parameters, which can be seen with 'crumble -h'. Some of these are the ones adjusted in the optimised crumble: -u30 adjusts the quality used in high confidence calls (defaults to 40); -Q60 reduces the minimum SNP consensus confidence required to trigger quality value replacement, from 70 (-9) or 75 (-1); likewise -D100 reduces the minimum indel consensus confidence, from -125 (-9) or 150 (-1).

The lightest compression level (crumble -1) is designed to cope better with subsequent remapping to different reference sequences, achieved by storing more lossless quality values in regions of low mapping score, potential collapsed repeats or missing insertions. However this requires a considerably larger amount of storage.

For the full 50x data set, to run `crumble -9p8` on chromosome 1 took 41 minutes elapsed time on a 2.2GHz Intel Xeon E5-2660, using 3Gb of RAM. Processing the entire genome (a 155Gb BAM file) took just over 10 hours, peaking at 3.8Gb of RAM.

The effect differs slightly per caller, although as expected the lowest level of lossy compression (crumble -1) was always closest to the original calls. Even so, crumble -1 gives a compressed quality size only 14% larger than the binary quantisation method using scores 4 and 28 introduced in the previous section. Crumble with the maximum optimised GATK parameters appears to also work well with bcftools and freebayes, indicating the optimisation is more related to the data rather than the caller.

Both higher levels of crumble tested give around 2.3 times better quality compression than the binary quantisation.

##### GATK HaplotypeCaller

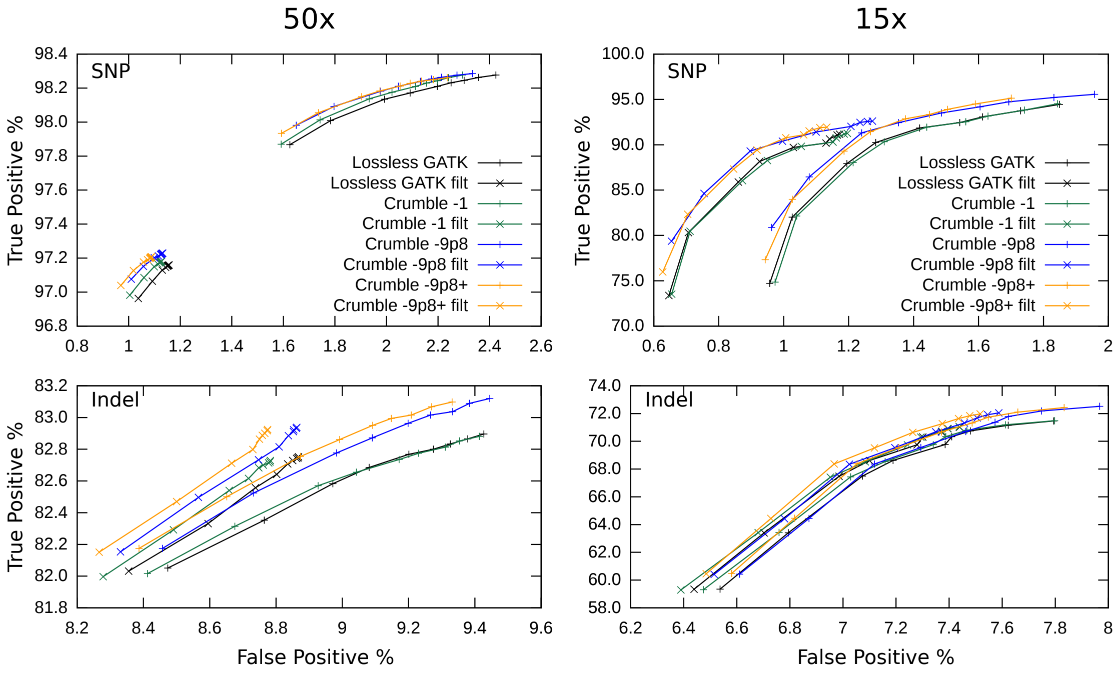


Figure 5: *True Positive vs False Negative rates of GATK HaplotypeCaller on the Crumbled vs lossless qualities.*

There is some variation between 50x / 15x and between SNP / Indel on whether the light Crumble -1 qualities are better than the lossless ones. However uniformly the P-score smoothing and more aggressive compression modes of Crumble are beneficial to all tests, with the more optimised parameters working best overall.

Table 19: GATK HC: 50x Crumble -1

| Type | | Q>0 | Q>=30 | Filtered |
|---|---|---|---|---|
| SNP | TP | 265007 | 264826 | 262030 |
| SNP | FP | 6226 | 5715 | 2968 |
| SNP | FN | 4648 | 4829 | 7625 |
| InDel | TP | 38155 | 38088 | 38064 |
| InDel | FP | 3965 | 3846 | 3649 |
| InDel | FN | 7881 | 7948 | 7972 |

**CRAM qual size 613,816,217**

Table 20: GATK HC: 15x Crumble -1

| Type | | Q>0 | Q>=30 | Filtered |
|---|---|---|---|---|
| SNP | TP | 254875 | 247918 | 242191 |
| SNP | FP | 4787 | 3624 | 2580 |
| SNP | FN | 14780 | 21737 | 27464 |
| InDel | TP | 32908 | 32116 | 32106 |
| InDel | FP | 2783 | 2544 | 2507 |
| InDel | FN | 13128 | 13920 | 13930 |

**CRAM qual size 260,305,104**

Table 21: GATK HC: 50x Crumble -9p8

| Type | | Q>0 | Q>=30 | Filtered |
|---|---|---|---|---|
| SNP | TP | 265032 | 264907 | 262161 |
| SNP | FP | 6334 | 5770 | 2980 |
| SNP | FN | 4623 | 4748 | 7494 |
| InDel | TP | 38265 | 38193 | 38157 |
| InDel | FP | 3991 | 3869 | 3699 |
| InDel | FN | 7771 | 7843 | 7879 |

**CRAM qual size 234,945,688**

Table 22: GATK HC: 15x Crumble -9p8

| Type | | Q>0 | Q>=30 | Filtered |
|---|---|---|---|---|
| SNP | TP | 257697 | 252166 | 246502 |
| SNP | FP | 5145 | 3804 | 2742 |
| SNP | FN | 11958 | 17489 | 23153 |
| InDel | TP | 33384 | 32549 | 32538 |
| InDel | FP | 2890 | 2625 | 2581 |
| InDel | FN | 12652 | 13487 | 13498 |

**CRAM qual size 77,416,003**

Table 23: GATK HC: 50x Crumble -9p8 -u30 -Q60 -D100

| Type | | Q>0 | Q>=30 | Filtered |
|---|---|---|---|---|
| SNP | TP | 264966 | 264834 | 262100 |
| SNP | FP | 6059 | 5551 | 2866 |
| SNP | FN | 4689 | 4821 | 7555 |
| InDel | TP | 38255 | 38187 | 38147 |
| InDel | FP | 3937 | 3819 | 3658 |
| InDel | FN | 7781 | 7849 | 7889 |

**CRAM qual size 228,658,529**

Table 24: GATK HC: 15x Crumble -9p8 -u30 -Q60 -D100

| Type | | Q>0 | Q>=30 | Filtered |
|---|---|---|---|---|
| SNP | TP | 256536 | 250405 | 244759 |
| SNP | FP | 4439 | 3491 | 2488 |
| SNP | FN | 13119 | 19250 | 24896 |
| InDel | TP | 33344 | 32534 | 32521 |
| InDel | FP | 2834 | 2589 | 2547 |
| InDel | FN | 12692 | 13502 | 13515 |

**CRAM qual size 72,072,237**

##### Bcftools

The affect of Crumble on bcftools is less clear than GATK, particularly at 50x. Not visible in the plot, the lossless and Crumble -1 SNP lines are superimposed for the 15x sample, possibly because at shallow data fewer quality values are adjusted. Which algorithm works best varies slightly based on which quality score is used in filtering, but the winner for SNPs is usually one of the two highest Crumble levels. Indels show less significant differences after filtering, perhaps due to lack of using quality in the filtering, with all 4 methods picking a slightly different trade off between precision and specificity.

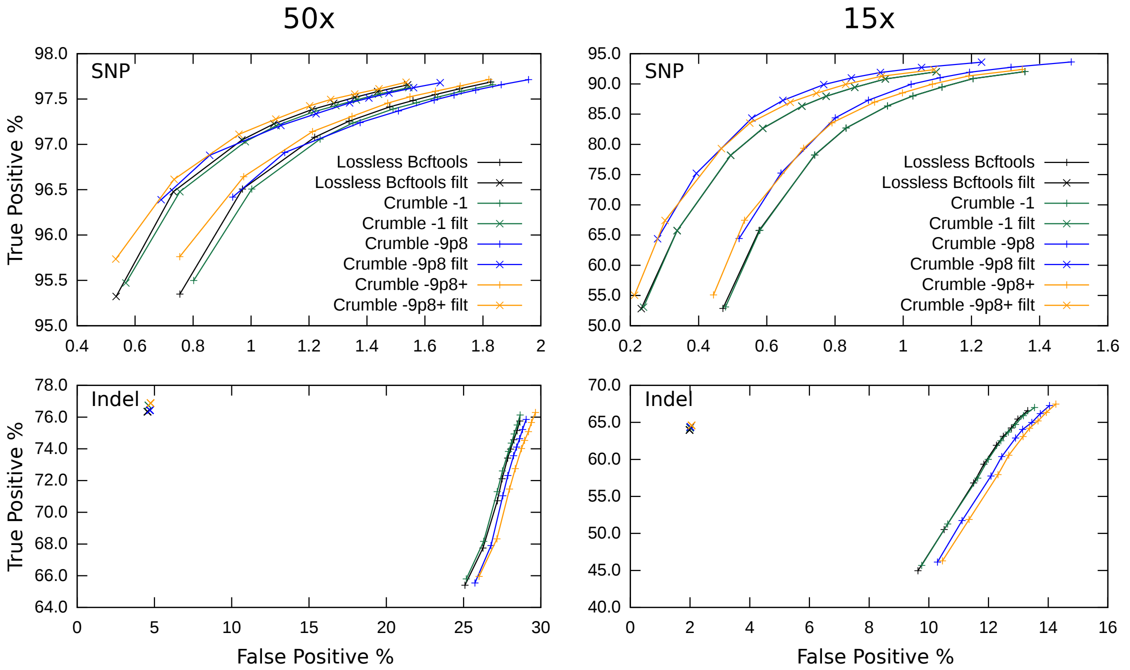


Figure 6: *True Positive vs False Negative rates of Bcftools on the Crumbled vs lossless qualities.*

Table 25: Bcftools: 50x crumble -1

| Type | | Q>0 | Q>=30 | Filtered |
|---|---|---|---|---|
| SNP | TP | 263659 | 262617 | 262534 |
| SNP | FP | 5496 | 3972 | 3234 |
| SNP | FN | 5996 | 7038 | 7121 |
| InDel | TP | 35618 | 33992 | 35327 |
| InDel | FP | 14561 | 13156 | 1710 |
| InDel | FN | 10418 | 12044 | 10709 |

**CRAM qual size 613,816,217**

Table 26: Bcftools: 15x Crumble -1

| Type | | Q>0 | Q>=30 | Filtered |
|---|---|---|---|---|
| SNP | TP | 253190 | 232872 | 232748 |
| SNP | FP | 4764 | 2243 | 1647 |
| SNP | FN | 16465 | 36783 | 36907 |
| InDel | TP | 31980 | 28783 | 29591 |
| InDel | FP | 5307 | 4079 | 605 |
| InDel | FN | 14056 | 17253 | 16445 |

**CRAM qual size 260,305,104**

Table 27: Bcftools: 50x crumble -9p8

| Type | | Q>0 | Q>=30 | Filtered |
|---|---|---|---|---|
| SNP | TP | 263766 | 262883 | 262798 |
| SNP | FP | 5818 | 4361 | 3569 |
| SNP | FN | 5889 | 6772 | 6857 |
| InDel | TP | 35469 | 33868 | 35186 |
| InDel | FP | 14801 | 13321 | 1740 |
| InDel | FN | 10567 | 12168 | 10850 |

**CRAM qual size 234,945,688**

Table 28: Bcftools: 15x Crumble -9p8

| Type | | Q>0 | Q>=30 | Filtered |
|---|---|---|---|---|
| SNP | TP | 256171 | 242505 | 242379 |
| SNP | FP | 5675 | 2507 | 1873 |
| SNP | FN | 13484 | 27150 | 27276 |
| InDel | TP | 32053 | 28951 | 29643 |
| InDel | FP | 5566 | 4291 | 608 |
| InDel | FN | 13983 | 17085 | 16393 |

**CRAM qual size 77,416,003**

Table 29: Bcftools: 50x crumble -9p8 -u30 -Q60 -D100

| Type | | Q>0 | Q>=30 | Filtered |
|---|---|---|---|---|
| SNP | TP | 263799 | 262793 | 262710 |
| SNP | FP | 5454 | 3925 | 3197 |
| SNP | FN | 5856 | 6862 | 6945 |
| InDel | TP | 35674 | 34073 | 35394 |
| InDel | FP | 15310 | 13747 | 1765 |
| InDel | FN | 10362 | 11963 | 10642 |

**CRAM qual size 228,658,529**

Table 30: Bcftools: 15x Crumble -9p8 -u30 -Q60 -D100

| Type | | Q>0 | Q>=30 | Filtered |
|---|---|---|---|---|
| SNP | TP | 253740 | 234599 | 234475 |
| SNP | FP | 4909 | 2169 | 1579 |
| SNP | FN | 15915 | 35056 | 35180 |
| InDel | TP | 32146 | 29044 | 29732 |
| InDel | FP | 5681 | 4400 | 623 |
| InDel | FN | 13890 | 16992 | 16304 |

**CRAM qual size 72,072,237**

##### Freebayes

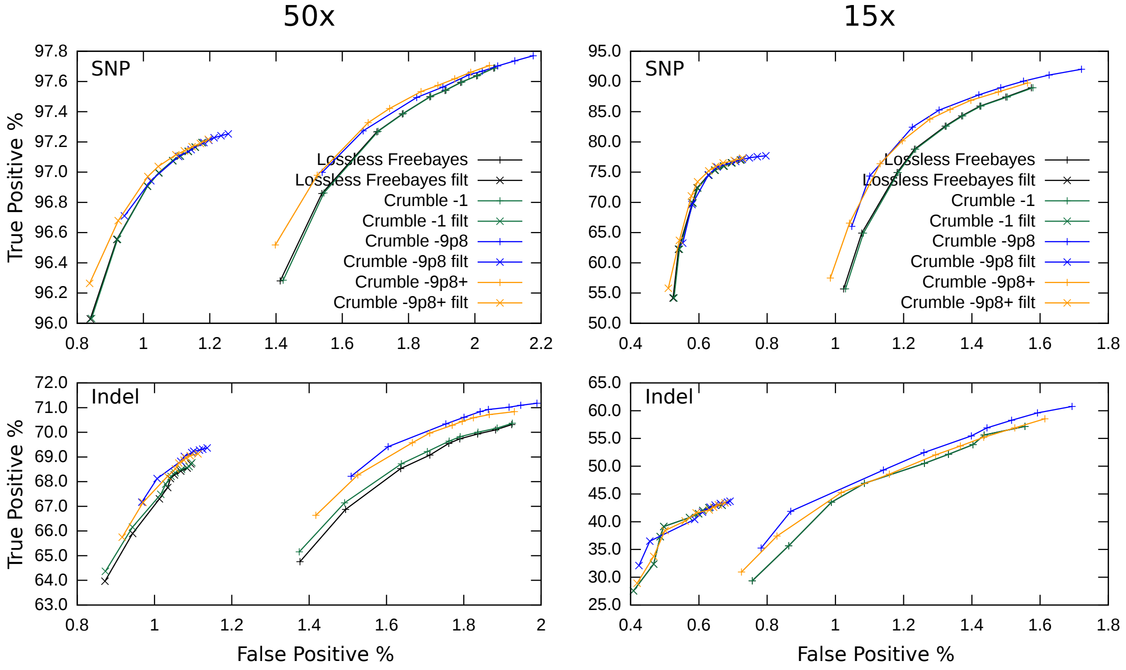


Figure 7: *True Positive vs False Negative rates of Freebayes on the Crumbled vs lossless qualities.*

With Freebayes, as with Bcftools, the lossless and Crumble -1 lines are superimposed. Crumble makes little difference to SNP calling after filtering, although there are slight gains with the GATK-optimised parameters. For indels after filtering the more compressed -9p8 options give a slight improvement at 50x.

Table 31: Freebayes: 50x crumble -1

| Type | | Q>0 | Q>=30 | Filtered |
|---|---|---|---|---|
| SNP | TP | 264319 | 262915 | 261772 |
| SNP | FP | 6026 | 5002 | 2881 |
| SNP | FN | 5336 | 6740 | 7883 |
| InDel | TP | 32759 | 32060 | 31403 |
| InDel | FP | 677 | 575 | 331 |
| InDel | FN | 13277 | 13976 | 14633 |

**CRAM qual size 613,816,217**

Table 32: Freebayes: 15x Crumble -1

| Type | | Q>0 | Q>=30 | Filtered |
|---|---|---|---|---|
| SNP | TP | 258868 | 222752 | 200889 |
| SNP | FP | 5014 | 2991 | 1273 |
| SNP | FN | 10787 | 46903 | 68766 |
| InDel | TP | 30122 | 23260 | 18760 |
| InDel | FP | 535 | 297 | 108 |
| InDel | FN | 15914 | 22776 | 27276 |

**CRAM qual size 260,305,104**

Table 33: Freebayes: 50x crumble -9p8

| Type | | Q>0 | Q>=30 | Filtered |
|---|---|---|---|---|
| SNP | TP | 264318 | 263302 | 262094 |
| SNP | FP | 6376 | 5324 | 3136 |
| SNP | FN | 5337 | 6353 | 7561 |
| InDel | TP | 32974 | 32610 | 31847 |
| InDel | FP | 716 | 612 | 353 |
| InDel | FN | 13062 | 13426 | 14189 |

**CRAM qual size 234,945,688**

Table 34: Freebayes: 15x Crumble -9p8

| Type | | Q>0 | Q>=30 | Filtered |
|---|---|---|---|---|
| SNP | TP | 258916 | 236683 | 207011 |
| SNP | FP | 6004 | 3410 | 1476 |
| SNP | FN | 10739 | 32972 | 62644 |
| InDel | TP | 30393 | 25528 | 19627 |
| InDel | FP | 597 | 362 | 125 |
| InDel | FN | 15643 | 20508 | 26409 |

**CRAM qual size 77,416,003**

Table 35: Freebayes: 50x crumble -9p8 -u30 -Q60 -D100

| **Type** | | **Q>0** | **Q>=30** | **Filtered** |
|---|---|---|---|---|
| SNP | TP | 264312 | 263002 | 261876 |
| SNP | FP | 5976 | 4923 | 2907 |
| SNP | FN | 5343 | 6653 | 7779 |
| InDel | TP | 32865 | 32357 | 31651 |
| InDel | FP | 689 | 583 | 340 |
| InDel | FN | 13171 | 13679 | 14385 |

**CRAM qual size 228,658,529**

Table 36: Freebayes: 15x Crumble -9p8 -u30 -Q60 -D100

| **Type** | | **Q>0** | **Q>=30** | **Filtered** |
|---|---|---|---|---|
| SNP | TP | 258853 | 225856 | 202815 |
| SNP | FP | 5065 | 2921 | 1283 |
| SNP | FN | 10802 | 43799 | 66840 |
| InDel | TP | 30189 | 23959 | 19150 |
| InDel | FP | 559 | 314 | 114 |
| InDel | FN | 15847 | 22077 | 26886 |

**CRAM qual size 72,072,237**

##### 4.3 CALQ

CALQ requires a sorted SAM file plus reference sequence as input and emits a new file containing the compressed qualities in its own format. The decode process produces a file containing just qualities, which with the aid of a supplied python script can then be put back into the original SAM file.

To encode:

```
calq -r $HREF -q Illumina-1.8+ -o CHM1_CHM13_2.chr1.sam.cq \
    -f CHM1_CHM13_2.chr1.sam 2>&1 | tee CHM1_CHM13_2.chr1.sam.calq.txt
```

To decode:

```
calq -f -s CHM1_CHM13_2.chr1.sam -d -o CHM1_CHM13_2.chr1.sam.cq.qual \
    CHM1_CHM13_2.chr1.sam.cq
```

Followed by `replace_qual_sam.py` to replace the qualities in the original input SAM file.

The encode process took approximately 7 hours for chromosome 1 and the decode 1.5 hours.

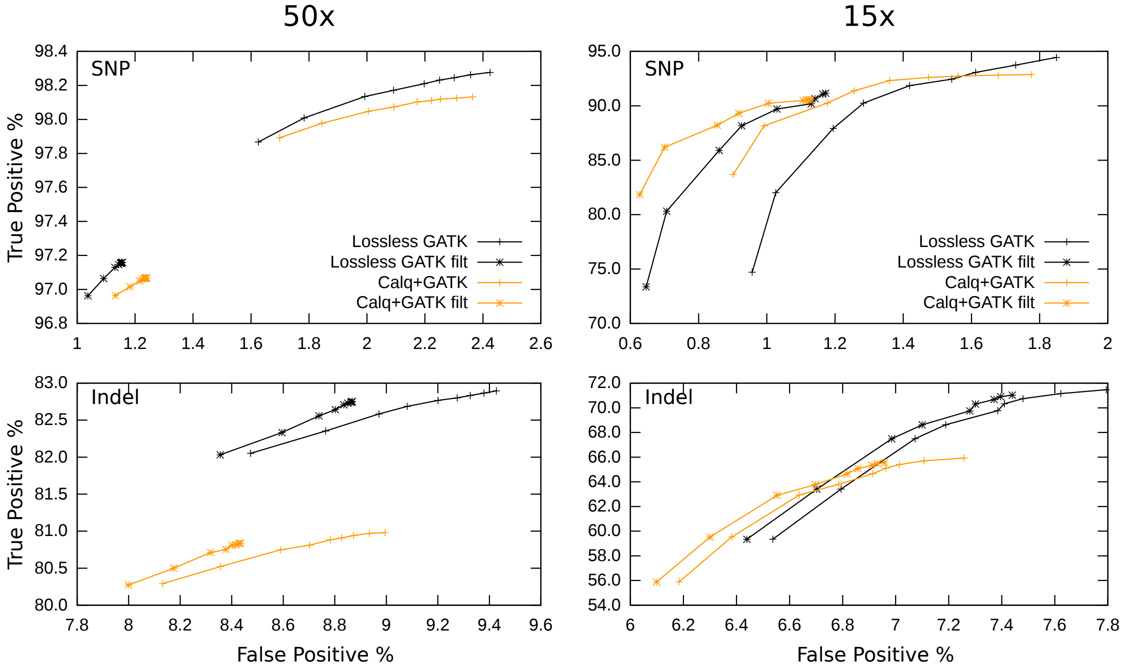


Figure 8: *True Positive vs False Negative rates of GATK HaplotypeCaller on the lossless vs CALQ qualities.*

We only show GATK HaplotypeCaller results for CALQ and QVZ2, as evaluating these tools is not the primary focus of this paper.

Compared to the lossless qualities, with the 50x data sets CALQ gives a significant decrease in true positives. The 15x data set fares better, representing a different tradeoff between precision and recall. The compressed quality size is comparable to the lightest compression with crumble ('crumble -1').

Table 37: CALQ + GATK HC, 50x

| **Type** | | **Q>0** | **Q>=30** | **Filtered** |
|---|---|---|---|---|
| SNP | TP | 264619 | 264539 | 261740 |
| SNP | FP | 6408 | 5877 | 3266 |
| SNP | FN | 5036 | 5116 | 7915 |
| InDel | TP | 37280 | 37235 | 37202 |
| InDel | FP | 3685 | 3585 | 3412 |
| InDel | FN | 8756 | 8801 | 8834 |

**CALQ .cq size 618,891,043**

Table 38: CALQ + GATK HC, 15x

| **Type** | | **Q>0** | **Q>=30** | **Filtered** |
|---|---|---|---|---|
| SNP | TP | 250452 | 248941 | 243309 |
| SNP | FP | 4527 | 3432 | 2469 |
| SNP | FN | 19203 | 20714 | 26346 |
| InDel | TP | 30348 | 29767 | 29761 |
| InDel | FP | 2375 | 2211 | 2177 |
| InDel | FN | 15688 | 16269 | 16275 |

**CALQ .cq size 187,994,047**

#### 4.4 QVZ2

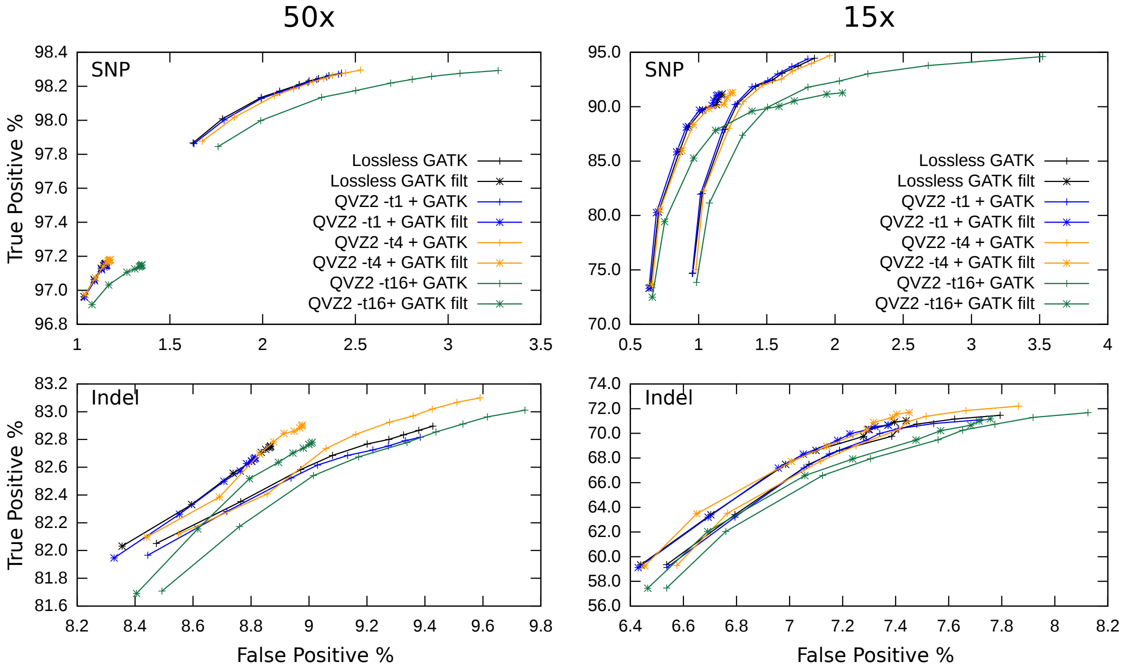


Figure 9: *True Positive vs False Negative rates of GATK HaplotypeCaller on the lossless vs QVZ2 qualities, 50x.*

QVZ2 operates on a file containing only quality values (e.g. every 4th line in a FASTQ file). It required around 10Gb of RAM and took 27 minutes to encode. It uses its own compressed file format for storing the quality values. After decoding we ran the `replace_qual_sam.py` tool from CALQ to update the SAM file prior to variant calling.

Comparing the Crumble results with QVZ2 we see the effect of minimising quality mean squared error vs aggressively increasing and decreasing qualities based on likelihood of variant calls changing. The mean squared error from Crumble changes will be very significant, but the size reduction is proportionally far greater while still achieving minimal changes to variant calling, in this case a small gain. QVZ2 has minimal impact on calling precision and recall at its lowest level (-t1). QVZ2 -t4 produces a slight shift towards more false positives with fewer false negatives, but is broadly beneficial, especially post filtering. The compression ratio at this option is not far behind CALQ and Crumble -1. Finally QVZ2 -t16 gives the smallest file of all (about 10% smaller than crumble -9p8), but has a significant increase in false positives.

Table 39: QVZ2 -t 1 + GATK HC, 50x

| **Type** | | **Q>0** | **Q>=30** | **Filtered** |
|---|---|---|---|---|
| SNP | TP | 264991 | 264810 | 261954 |
| SNP | FP | 6541 | 5947 | 3052 |
| SNP | FN | 4664 | 4845 | 7701 |
| InDel | TP | 38125 | 38065 | 38038 |
| InDel | FP | 3948 | 3826 | 3663 |
| InDel | FN | 7911 | 7971 | 7998 |

**QVZ2 qual size: 1,493,843,021**

Table 40: QVZ2 -t 1 + GATK HC, 15x

| **Type** | | **Q>0** | **Q>=30** | **Filtered** |
|---|---|---|---|---|
| SNP | TP | 254503 | 247598 | 241799 |
| SNP | FP | 4668 | 3495 | 2457 |
| SNP | FN | 15152 | 22057 | 27856 |
| InDel | TP | 32732 | 31970 | 31964 |
| InDel | FP | 2739 | 2514 | 2473 |
| InDel | FN | 13304 | 14066 | 14072 |

**QVZ2 qual size: 441,580,609**

Table 41: QVZ2 -t 4 + GATK HC, 50x

| Type | | Q>0 | Q>=30 | Filtered |
|---|---|---|---|---|
| SNP | TP | 265058 | 264873 | 262025 |
| SNP | FP | 6874 | 6155 | 3095 |
| SNP | FN | 4597 | 4782 | 7630 |
| InDel | TP | 38256 | 38175 | 38138 |
| InDel | FP | 4058 | 3904 | 3732 |
| InDel | FN | 7780 | 7861 | 7898 |

**QVZ2 qual size: 657,068,110**

Table 42: QVZ2 -t 4 + GATK HC, 15x

| Type | | Q>0 | Q>=30 | Filtered |
|---|---|---|---|---|
| SNP | TP | 255342 | 248070 | 242197 |
| SNP | FP | 5105 | 3706 | 2635 |
| SNP | FN | 14313 | 21585 | 27458 |
| InDel | TP | 33240 | 32373 | 32365 |
| InDel | FP | 2837 | 2592 | 2557 |
| InDel | FN | 12796 | 13663 | 13671 |

**QVZ2 qual size: 194,172,554**

Table 43: QVZ2 -t 16 + GATK HC, 50x

| Type | | Q>0 | Q>=30 | Filtered |
|---|---|---|---|---|
| SNP | TP | 265051 | 264852 | 261936 |
| SNP | FP | 8959 | 7322 | 3545 |
| SNP | FN | 4604 | 4803 | 7719 |
| InDel | TP | 38215 | 38108 | 38073 |
| InDel | FP | 4126 | 3925 | 3740 |
| InDel | FN | 7821 | 7928 | 7963 |

**QVZ2 qual size: 201,725,874**

Table 44: QVZ2 -t 16 + GATK HC, 15x

| Type | | Q>0 | Q>=30 | Filtered |
|---|---|---|---|---|
| SNP | TP | 255105 | 247508 | 241601 |
| SNP | FP | 9314 | 4541 | 3410 |
| SNP | FN | 14550 | 22147 | 28054 |
| InDel | TP | 32997 | 31986 | 31977 |
| InDel | FP | 2918 | 2616 | 2584 |
| InDel | FN | 13039 | 14050 | 14059 |

**QVZ2 qual size: 59,859,656**

#### 4.5 Syndip regions

The Syndip data set is not perfect and there are bed files to filter out poor regions. This may lead to concern that we are testing only well behaved data and do not know how the tools work in hard to sequence regions. This concern is true for all truth sets generated from real sequencing data, including the Genome in a Bottle (GIAB) and Platinum Genomes (PlatGen) data sets that have been established for longer. The Syndip paper indicates that testing variant callers on Syndip probes more of the genome, including more difficult parts, leading to substantially higher false positive rates than seen with GIAB and PlatGen.

> "*Figure 2a reveals that the FPPM of SNPs estimated from Syndip is often 5-10 times higher than FPPM estimated from GIAB or PlatGen. Looking into the Syndip FP SNPs, we found most of them are located in CNVs that are evident in PacBio data in the context of long flanking regions, but look dubious in short-read data alone.*"

The total number of bases included in chromosome 1 from Syndip is 212.9Mb out of 225.3Mb of non-N reference. This compares favourably to 204.4Mb in filtered GIAB.

Furthermore we can subtract the GIAB regions from Syndip regions to get only regions that occur in Syndip (around 8.4Mb). To see a significantly elevated overall false positive rate, either the Syndip data is highly erroneous or the bulk of the extra false positives are within this region. To test this we ran GATK on the data sets filtered to this region alone.

Comparing this to the full Syndip regions for chromosome 1 we see that 65% of false positives occur within this small portion. This addresses the possibility that we are restricting ourselves to only good quality data. The results show that Crumble still performs well in this region.

Table 45: GATK HC: 50x Original

| Type | | Q>0 | Q>=30 | Filtered |
|---|---|---|---|---|
| SNP | TP | 19858 | 19740 | 18298 |
| SNP | FP | 4384 | 4014 | 1950 |
| SNP | FN | 3818 | 3936 | 5378 |
| InDel | TP | 9822 | 9774 | 9752 |
| InDel | FP | 2992 | 2913 | 2807 |
| InDel | FN | 4416 | 4464 | 4486 |

Table 46: GATK HC: 15x Original

| Type | | Q>0 | Q>=30 | Filtered |
|---|---|---|---|---|
| SNP | TP | 17411 | 16711 | 15616 |
| SNP | FP | 3063 | 2553 | 1643 |
| SNP | FN | 6265 | 6965 | 8060 |
| InDel | TP | 7187 | 6966 | 6961 |
| InDel | FP | 2040 | 1923 | 1887 |
| InDel | FN | 7051 | 7272 | 7277 |

Table 47: GATK HC: 50x Crumble -9p8...

| Type | | Q>0 | Q>=30 | Filtered |
|---|---|---|---|---|
| SNP | TP | 19900 | 19801 | 18360 |
| SNP | FP | 4229 | 3872 | 1829 |
| SNP | FN | 3776 | 3875 | 5316 |
| InDel | TP | 9879 | 9822 | 9789 |
| InDel | FP | 2985 | 2902 | 2792 |
| InDel | FN | 4359 | 4416 | 4449 |

Table 48: GATK HC: 15x Crumble -9p8...

| Type | | Q>0 | Q>=30 | Filtered |
|---|---|---|---|---|
| SNP | TP | 17711 | 16990 | 15899 |
| SNP | FP | 2948 | 2496 | 1624 |
| SNP | FN | 5965 | 6686 | 7777 |
| InDel | TP | 7399 | 7130 | 7125 |
| InDel | FP | 2080 | 1921 | 1884 |
| InDel | FN | 6839 | 7108 | 7113 |

Table 49: GATK HC: 50x Calq

| Type | | Q>0 | Q>=30 | Filtered |
|---|---|---|---|---|
| SNP | TP | 19521 | 19465 | 18050 |
| SNP | FP | 4203 | 3931 | 1999 |
| SNP | FN | 4155 | 4211 | 5626 |
| InDel | TP | 9186 | 9153 | 9123 |
| InDel | FP | 2751 | 2688 | 2576 |
| InDel | FN | 5052 | 5085 | 5115 |

Table 50: GATK HC: 15x Calq

| Type | | Q>0 | Q>=30 | Filtered |
|---|---|---|---|---|
| SNP | TP | 16267 | 16065 | 15020 |
| SNP | FP | 2726 | 2367 | 1525 |
| SNP | FN | 7409 | 7611 | 8656 |
| InDel | TP | 6042 | 5913 | 5908 |
| InDel | FP | 1716 | 1628 | 1598 |
| InDel | FN | 8196 | 8325 | 8330 |

Table 51: GATK HC: 50x QVZ2 -t 4

| Type | | Q>0 | Q>=30 | Filtered |
|---|---|---|---|---|
| SNP | TP | 19901 | 19771 | 18335 |
| SNP | FP | 4486 | 4087 | 1981 |
| SNP | FN | 3775 | 3905 | 5341 |
| InDel | TP | 9894 | 9829 | 9799 |
| InDel | FP | 3038 | 2932 | 2833 |
| InDel | FN | 4344 | 4409 | 4439 |

Table 52: GATK HC: 15x QVZ2 -t 4

| Type | | Q>0 | Q>=30 | Filtered |
|---|---|---|---|---|
| SNP | TP | 17588 | 16842 | 15729 |
| SNP | FP | 3234 | 2652 | 1730 |
| SNP | FN | 6088 | 6834 | 7947 |
| InDel | TP | 7358 | 7097 | 7083 |
| InDel | FP | 2078 | 1940 | 1900 |
| InDel | FN | 6880 | 7141 | 7155 |

### 5 Syndip Summary

Table 53: Summary of filtered 50x, Syndip Chromosome 1

| Tool | Method | SNP | | Indel | | Size |
|---|---|---|---|---|---|---|
| | | FP | FN | FP | FN | |
| GATK | Lossless | 3047 | 7678 | 3690 | 7961 | 4,106,563,351 |
| GATK | Qual 4 + 28 | 2950 | 8010 | 3377 | 8798 | 539,249,433 |
| GATK | Qual 25 | 3189 | 8360 | 3402 | 8721 | 756,507 |
| GATK | Crumble -1 | 2968 | 7625 | 3649 | 7972 | 613,816,217 |
| GATK | Crumble -9p8 | 2980 | 7494 | 3699 | 7879 | 234,945,688 |
| GATK | Crumble -9p8 -u30... | 2866 | 7555 | 3658 | 7889 | 228,658,529 |
| GATK | CALQ | 3266 | 7915 | 3412 | 8834 | 618,891,043 |
| GATK | QVZ2 -t1 | 3052 | 7701 | 3663 | 7998 | 1,493,843,021 |
| GATK | QVZ2 -t4 | 3095 | 7630 | 3732 | 7898 | 657,068,110 |
| GATK | QVZ2 -t16 | 3545 | 7719 | 3740 | 7963 | 201,725,874 |
| Bcftools | Lossless | 3216 | 7056 | 1678 | 10893 | 4,106,563,351 |
| Bcftools | Qual 4 + 28 | 3171 | 7148 | 1652 | 10956 | 539,249,433 |
| Bcftools | Qual 25 | 4515 | 7116 | 1567 | 11472 | 756,507 |
| Bcftools | Crumble -1 | 3234 | 7121 | 1710 | 10709 | 613,816,217 |
| Bcftools | Crumble -9p8 | 3569 | 6857 | 1740 | 10850 | 234,945,688 |
| Bcftools | Crumble -9p8 -u30... | 3197 | 6945 | 1765 | 10642 | 228,658,529 |
| Freebayes | Lossless | 2880 | 7886 | 330 | 14674 | 4,106,563,351 |
| Freebayes | Qual 4 + 28 | 2789 | 8018 | 324 | 14877 | 539,249,433 |
| Freebayes | Qual 25 | 4147 | 7833 | 330 | 14239 | 756,507 |
| Freebayes | Crumble -1 | 2881 | 7883 | 331 | 14633 | 613,816,217 |
| Freebayes | Crumble -9p8 | 3136 | 7561 | 353 | 14189 | 234,945,688 |
| Freebayes | Crumble -9p8 -u30... | 2907 | 7779 | 340 | 14385 | 228,658,529 |

Table 54: Summary of filtered 15x, Syndip Chromosome 1

| Tool | Method | SNP | | Indel | | Size |
|---|---|---|---|---|---|---|
| | | FP | FN | FP | FN | |
| GATK | Lossless | 2517 | 27761 | 2521 | 13925 | 1,211,486,517 |
| GATK | Qual 4 + 28 | 2206 | 31382 | 2133 | 16152 | 159,104,061 |
| GATK | Qual 25 | 3132 | 32732 | 2236 | 15585 | 223,176 |
| GATK | Crumble -1 | 2580 | 27464 | 2507 | 13930 | 260,305,104 |
| GATK | Crumble -9p8 | 2742 | 23153 | 2581 | 13498 | 77,416,003 |
| GATK | Crumble -9p8 -u30... | 2488 | 24896 | 2547 | 13515 | 72,072,237 |
| GATK | CALQ | 2469 | 26346 | 2177 | 16275 | 187,994,047 |
| GATK | QVZ2 -t1 | 2457 | 27856 | 2473 | 14072 | 441,580,609 |
| GATK | QVZ2 -t4 | 2635 | 27458 | 2557 | 13671 | 194,172,554 |
| GATK | QVZ2 -t16 | 3410 | 28054 | 2584 | 14059 | 59,859,656 |
| Bcftools | Lossless | 1648 | 36921 | 596 | 16586 | 1,211,486,517 |
| Bcftools | Qual 4 + 28 | 1613 | 38738 | 594 | 16682 | 159,104,061 |
| Bcftools | Qual 25 | 2088 | 40804 | 557 | 17254 | 223,176 |
| Bcftools | Crumble -1 | 1647 | 36907 | 605 | 16445 | 260,305,104 |
| Bcftools | Crumble -9p8 | 1873 | 27276 | 608 | 16393 | 77,416,003 |
| Bcftools | Crumble -9p8 -u30... | 1579 | 35180 | 623 | 16304 | 72,072,237 |
| Freebayes | Lossless | 1269 | 68763 | 108 | 27276 | 1,211,486,517 |
| Freebayes | Qual 4 + 28 | 1236 | 70514 | 99 | 27574 | 159,104,061 |
| Freebayes | Qual 25 | 1455 | 69174 | 118 | 26851 | 223,176 |
| Freebayes | Crumble -1 | 1273 | 68766 | 108 | 27276 | 260,305,104 |
| Freebayes | Crumble -9p8 | 1476 | 62644 | 125 | 26409 | 77,416,003 |
| Freebayes | Crumble -9p8 -u30... | 1283 | 66840 | 114 | 26886 | 72,072,237 |

### 6 Further compression

Unlike QVZ2 and CALQ, Crumble does not output compressed qualities itself. It is designed to be used in conjunction with an existing file format, ideally one that has efficient encoding of quality values. This means it works well in conjunction with CRAM, but improving compressibility of qualities also helps BAM.

The 15x sub-sampled file with and without Crumble for the single chromosome 1 test above have the following sizes:

| file | bytes |
|---|---|
| CHM1_CHM13_2.15x.chr1.bam | 3963702044 |
| CHM1_CHM13_2.15x.chr1.cram | 2188724919 |
| CHM1_CHM13_2.15x.chr1.crumble-opt.bam | 2325189762 |
| CHM1_CHM13_2.15x.chr1.crumble-opt.cram | 1049588799 |

In absolute bytes saved, BAM reduces by more (1.6 vs 1.1 Gb), due to initially poor compression of qualities. However in ratio terms, the original lossless CRAM was 45% smaller than the original BAM, but after Crumble the lossy CRAM is now 55% smaller than the corresponding BAM.

This particular data set has been through the GATK Base Quality Score Recalibration (BQSR) process which has preserved original qualities in the SAM OQ:Z tag. The `cram_size` tool from the Staden *io_lib* package gives summaries of the space taken by each data type within a CRAM file. The original and crumbled version are shown below for chromosome 1 of the 15x Syndip data set along with annotation of the most significant SAM fields.

```
Block content_id      11, total size   147342810 g     RN   (read names)
Block content_id      12, total size  1211486517     R QS   (quality scores)
Block content_id      13, total size      210086 g     IN   (bases in insertions)
Block content_id      14, total size    31483343    rR SC   (bases in soft-clips)
Block content_id      15, total size     7866518     R BF   (BAM flags)
Block content_id      16, total size     3517731    rR CF   (CRAM flags)
Block content_id      17, total size    13906529 g  r  AP   (POS field)
Block content_id      18, total size    13921662    r  RG   (Read group)
Block content_id      19, total size     1900911 g  r  MQ   (Mapping quality)
Block content_id      20, total size      355913 g  r  NS   (Mate reference ID)
Block content_id      21, total size      384498    r  MF   (Mate flags)
Block content_id      22, total size     2811406 g     TS   (TLEN field)
Block content_id      23, total size     5262570 g     NP   (PNEXT field)
Block content_id      24, total size     7926491 g     NF   (Read pairing)
Block content_id      26, total size     7764331    r  FN   (Feature (diff) count)
Block content_id      27, total size     2999582    rR FC   (Feature code)
Block content_id      28, total size    35781940 g  r  FP   (Feature position)
Block content_id      29, total size      155914 g  r  DL   (Length of CIGAR "D")
Block content_id      30, total size     5926103    rR BA   (Bases)
Block content_id      31, total size     8649685    rR BS   (Base substitions)
Block content_id      32, total size     3087067    r  TL   (Aux. tag list)
Block content_id 4281155, total size     6309393    r  ASC  (AS:i: aux tag)
Block content_id 4281187, total size     3410458 g     ASc  (AS:i: aux tag)
Block content_id 5063514, total size    14956889 g     MCZ  (MC:Z: aux tag)
Block content_id 5063770, total size         686 g     MDZ  (MD:Z: aux tag)
Block content_id 5067107, total size     2031763 g  r  MQc  (MQ:i: aux tag)
Block content_id 5131619, total size          66 g     NMc  (NM:i: aux tag)
Block content_id 5194586, total size      155949 g     OCZ  (OC:Z: aux tag)
Block content_id 5197929, total size       42528 g     OPi  (OP:i: aux tag)
Block content_id 5198170, total size   601811789     R OQZ  (OQ:Z: aux tag)
Block content_id 5261146, total size    29615589 g     PGZ  (PG:Z: aux tag)
Block content_id 5456218, total size     2021083 g     SAZ  (SA:Z: aux tag)
Block content_id 5591363, total size      602922 g     UQC  (UQ:i: aux tag)
Block content_id 5591395, total size    11069115    r  UQc  (UQ:i: aux tag)
Block content_id 5591411, total size      289324 g     UQs  (UQ:i: aux tag)
Block content_id 5787235, total size          42 g     XNc  (XN:i: aux tag)
Block content_id 5788739, total size      141166 g     XTC  (XT:i: aux tag)
```

```
Block content_id 5788771, total size       211183 g     XTc  (XT:i: aux tag)
```

Crumbled: as above, but with QS (quality scores) data series as:

```
Block content_id       12, total size    72072237     R QS
```

After this the next largest blocks are the original qualities (OQZ) as output as part of GATK BQSR and read query names (RN).

The original qualities can be completely discarded, as is now the recommendation in the GATK best practices. The other large auxiliary tag we safely remove is PG, as in this particular data it is both superfluous (existing only to inform which subset in a map-reduce style processing pipeline the read came from) and unfortunately also incorrect (none of the per-read PG tags match the @PG SAM header lines).

When all reads from the same template occur within the same CRAM slice the read names may be discarded without affecting variant calling and without losing pairing information as this is held in the CRAM NF data series. Long distance read pairs have their names retained to ensure pairing information is kept intact.

Crumble supports removal of both read names and specific auxiliary tags, as illustrated in the command below:

```
crumble -T OQ,PG -O cram,lossy_names -9p8 -u30 -Q60 -D100 \
  CHM1_CHM13_2.15x.chr1.cram CHM1_CHM13_2.15x.chr1.crumble-opt.cram
```

The CRAM file now has no OQ:Z or PG:Z blocks and read names consume 15,480,044 bytes instead of 147,342,810.

Repeating this test on the whole genome, at full depth (50x) and reduced depths of 30x and 15x, yields the file sizes show below. Comparison between BAM and CRAM sizes show that the benefits of using a columnar storage are significantly greater on the crumbled data.

| file | BAM bytes | CRAM bytes |
|---|---:|---:|
| CHM1_CHM13_2.all.lossless | 165,881,395,078 | 94,722,033,125 |
| CHM1_CHM13_2.all.crumble-opt | 42,971,979,964 | 12,735,423,262 |
| CHM1_CHM13_2.30x.lossless | 100,407,390,797 | 56,798,653,417 |
| CHM1_CHM13_2.30x.crumble-opt | 26,826,816,872 | 7,635,898,756 |
| CHM1_CHM13_2.15x.lossless | 51,338,937,983 | 28,416,618,181 |
| CHM1_CHM13_2.15x.crumble-opt | 14,483,060,883 | 3,862,050,827 |

An approximate breakdown of storage in the reduced CRAM for the complete 30x sample is 24% qualities, 16% remaining auxiliary tags, 12% soft-clipped bases, 5% remaining read names, 5% read groups, 4% alignment position and the remaining 34% alignment and sequence-reference differences plus a small amount of overhead.

#### 7 Other data sets

Although no truth sets are used for evaluating variation calling, we ran crumble on a variety of other data sets to report the size reduction when using `crumble -O cram,lossy_names -9p8`. The output was then converted back to BAM to compare the file size between formats.

Data sets chosen were a 420x deep E.Coli Illumina MiSeq run (MiSeq_Ecoli_DH10B_110721_PF) and an Illumina human RNASeq run (K562_cytosol_LID8465_TopHat_v2). These were taken from the Moving Picture Experts Group (MPEG, JTC1/SC29/WG11 committee) data set for on-going development of the MPEG-G format.

See `https://github.com/sfu-compbio/compression-benchmark/blob/master/samples.md` for download links. We avoided Oxford Nanopore Technology and Pacific Biosciences data as Crumble has not been evaluated on these yet.

Before and after file sizes are reported along with the space taken up by quality values and read names where applicable.

| File | Format | Method | Total size | Quality size | Name size |
|---|---|---|---|---|---|
| MiSeq_Ecoli_DH10B_110721_PF | BAM | Original | 1411850544 | n/a | n/a |
| MiSeq_Ecoli_DH10B_110721_PF | CRAM | Original | 862693214 | 714245853 | 65303357 |
| MiSeq_Ecoli_DH10B_110721_PF | BAM | Crumble -9p8 | 382629759 | n/a | n/a |
| MiSeq_Ecoli_DH10B_110721_PF | CRAM | Crumble -9p8 | 110838273 | 22526180 | 5197781 |
| K562_cytosol_LID8465_TopHat_v2 | BAM | Original | 13756734292 | n/a | n/a |
| K562_cytosol_LID8465_TopHat_v2 | CRAM | Original | 9323049595 | 6443896054 | 1655419459 |
| K562_cytosol_LID8465_TopHat_v2 | BAM | Crumble -9p8 | 4625068006 | n/a | n/a |
| K562_cytosol_LID8465_TopHat_v2 | CRAM | Crumble -9p8 | 2736614367 | 417779131 | 1085784399 |

The effect of Crumble on both data sets is a considerable reduction to quality size within CRAM (32 fold and 15 fold respectively). The ability to perform lossy read name compression (which is part of CRAM rather than Crumble) is hampered on the RNASeq data by having reads split over larger regions and not colocating within the same CRAM slice and very few being labelled as properly paired. As a consequence the read names are the largest data type in the crumbled RNASeq data set. Neither of these files have an excessively large collection of auxiliary tags.

For both files the ratio of original to crumbled size is higher with CRAM (7.8 and 3.4) than BAM (3.7 and 3.0), demonstrating the benefit of combining lossy quality encoding with a columnar file format.

During preparation of this manuscript a bug was fixed affecting the speed of Crumble on RNAseq data. Thus the RNASeq TopHat data was processed using a more recent git commit (v0.8-4-g556c716). Both version 0.8 and 0.8-4 were tested on the E.Coli data and observed to give identical results. Final timings were 6 min 43 seconds for the E.Coli data and 599 min 11 seconds for the RNASeq data corresponding to unthreaded BAM processing speeds of 3.3Mb/s and 0.34 Mb/s, demonstrating that there is still some degraded CPU performance operating on RNASeq data sets.

#### 8 Conclusion

As expected, the 15x sample has fewer confident consensus bases than the 50x sample leading to a slightly lower quality compression ratio, however even at 15x there is sufficient confidence in calling to discard most quality values.

The original CRAM file for the 15x chromosome 1 comprised 24 million reads and 36 billion base pairs, giving 2.67 bits per lossless quality value. After optimal Crumble parameters were applied, this reduced to 0.16 bits per quality.

It is clear there are a lot of parameters that can be adjusted for controlling when to adjust quality values, and to which values. We have not exhaustively explored this search space. There are also open questions on the performance of Crumble on somatic / non-clonal samples, such as cancers, or mixed sample data sets. Hence we do not recommend the use of Crumble on such data without prior evaluation.

We also do not recommend usage of Crumble on non-Illumina data sets until further evaluation has been made.
